# A Pilot Study Using AI for Genetic Effects on Personality: ABO Blood Type in Japan/Korea

**DOI:** 10.1101/2023.07.15.549129

**Authors:** Masayuki Kanazawa

## Abstract

It is estimated that genetic factors contribute to approximately 50% of an individual’s personality. However, despite this estimate, statistical analyses have yet to establish a significant and consistent relationship between personality tests and genetic factors. Conversely, a large number of individuals in Japan, South Korea, and Taiwan believe in a connection between genetically determined ABO blood type and personality traits. This pilot study aimed to investigate this relationship by analyzing data from a Japanese survey (N=1,827) and a South Korean survey (N=482) using a combination of traditional statistical methods and AI. The findings demonstrated a relationship between ABO blood type and self-reported personality traits, as assessed through several single-question items, consistent with the anticipated outcomes. The findings of this study imply that the relationship between blood type and personality may extend the ABO blood type to other heritable personality traits. Additionally, this study also explores the influence of the level of interest on personality and the potential role of evolutionary psychology.

## 1. Introduction

### 1.1 Influence of Heredity Based on Twin Studies

The impact of heredity on personality traits is estimated to approximately 50% based on empirical investigations, including twin studies [1-4]. Twins can be categorized into two distinct types: monozygotic and dizygotic. Monozygotic twins originate from fertilized eggs that undergo subsequent separation during embryonic development, resulting in individuals with indistinguishable DNA sequences. Conversely, dizygotic twins possess genetic dissimilarities comparable to those observed among non-twin siblings, as each originates from an independent fertilization event involving a different sperm and egg. In humans, the chromosomal makeup is diploid, characterized by 23 pairs of chromosomes, totaling 46 in number. During the production of reproductive cells (sperms and eggs) in the parents, meiosis randomly selects one chromosome from each pair. Consequently, when chromosomes segregate, recombination occurs randomly within a subset of paired chromosomes, resulting in the fertilized egg inheriting 50% of its DNA from the father and 50% from the mother. Thus, while monozygotic twins exhibit strikingly similar DNA profiles, dizygotic twins, on average, share only half of their genetic makeup. Remarkably, monozygotic twins often display congruent physical attributes, such as hair color and facial features, whereas dizygotic twins exhibit less resemblance in these regards. Thus, monozygotic twins share the same gender, but dizygotic twins can differ. Note that recent research revealed that a few rare mutations arising during early embryogenesis make the DNA profiles of monozygotic twins not exactly identical [5-6].

These fundamental aspects have prompted investigations into the influence of heredity on various psychological and physical characteristics, including intelligence, academic achievement, personality traits, and mental illness. Statistical analyses comparing monozygotic and dizygotic twins have facilitated the exploration of the effects of genetic factors by considering the differences between these groups. Typically, parents raise both monozygotic and dizygotic twins in a similar manner, irrespective of their zygosity. Consequently, if psychological or physical traits were unaffected by genetic influences, the observed differences would primarily stem from postnatal experiences. Thus, the dissimilarities between monozygotic and dizygotic twins should be, on average, comparable. However, research findings have consistently demonstrated significantly smaller differences between monozygotic twins than dizygotic twins. These results underscore the partial genetic determination of psychological and physical characteristics. Previous investigations have estimated that heredity accounts for about 50% of the variations found in intelligence and personality, more than 90% in fingerprints, height, and weight, and 80% in certain mental disorders such as schizophrenia, autism, and ADHD [7-8].

### 1.2 Genome-wide Association Studies

Advancements in the field of life sciences have been remarkable in recent years, owing to technological breakthroughs such as the next-generation sequencers. Notably, in 2003, the International Human Genome Sequencing Consortium declared the successful decoding of the entire human genome. Subsequently, genome sequencing initiatives were initiated worldwide, including the pioneering 1000 Genomes Project launched in 2008. Presently, numerous commercial genome analysis services have emerged, enabling the direct investigation of the relationship between genomes and personality traits and abilities, which was previously unattainable through conventional methods. This is achieved through the application of mathematical approaches such as linear algebra to analyze genome data.

During the early 2000s, genome-wide association study (GWAS) emerged as a prominent method for investigating the impact of single nucleotide polymorphisms (SNPs), representing genetic variations in genomic information among individuals, and to what extent they influence phenotypes. Prior to GWAS, genetic analysis methods were primarily focused on monogenic traits, examining the effect of a single gene on a phenotype. However, it became evident that a single gene could not explain many phenotypic variations. Consequently, even when SNPs were identified through GWAS, the genetic contribution derived from them was typically considerably smaller than the estimates obtained from twin studies. This phenomenon, known as missing or hidden heritability, garnered large attention [2].

Nonetheless, the situation has undergone substantial change since approximately 2016. The discovery of over 1,200 SNPs that appeared to influence educational background marked a significant breakthrough. The estimated impact of these SNPs accounted for 12% of the heritability of intelligence [9]. In a subsequent paper published in 2022, utilizing a sample size of 3 million individuals, the explanatory power increased to 16 percent [10]. Similarly, an analysis of a dataset comprising 5 million individuals, predominantly of European descent, demonstrated the involvement of approximately SNPs in determining height [11]. Furthermore, it was observed that these findings could account for 40% of the inter-individual height differences among individuals of European ancestry.

### 1.3 History of Blood Type and Personality Studies

However, the current situation concerning personality has shown limited improvement. A notable illustration of this can be found in the study conducted by Lo et al. in 2017, which investigated the association between data from all SNPs in the human genome of over 260,000 individuals and the widely used the Big Five personality tests in psychology [12]. Their findings identified 6 genes associated with personality traits. Nevertheless, the coefficient of determination (*R*^*2*^), which indicates the degree of influence, was less than 0.4% for all identified genes, rendering them insignificantly small and typically regarded as errors. This is in stark contrast to the 50% figure obtained from the aforementioned twin studies. Furthermore, a study by Hindley et al. in 2023 improved the method to measure effects on multiple genes and expanded the gene pool to more than 400, resulting in an increase in *R*^*2*^, but in the lower 2% range [13].

The underlying factors contributing to this exceedingly small heritability in personality include: 1) interactions between genes involving nonlinear dynamics, 2) genetic and environmental interactions, and 3) difficulties associated with detecting and analyzing SNPs using current molecular genetic techniques. These factors bear significance in gaining a comprehensive understanding of genetic phenomena. They suggest that personality fundamentally operates as a complex system with intricate interactions between genetic predispositions and external influences, such as sociocultural factors.

Conversely, certain studies suggest that the influence of individual genes may be greater than that obtained by GWAS. For instance, an investigation explored the connection between ABO blood type (hereafter referred to as blood type) and personality, a subject that has been extensively studied in Japan and other East Asian countries for nearly a century [14-15]. In fact, approximately half of the population in Japan, South Korea (hereafter referred to as Korea), and Taiwan perceives a relationship between blood type and personality [16-18]. The prevalence of blood type knowledge due to military drafts and blood donation systems has led to widespread participation in this topic, not only by academics but also by independent researchers [19-20]. However, despite these efforts, a scientific consensus on the association between blood type and personality has yet to be reached [21-22].

An important commonality among previous studies is the utilization of multiple-question items, up to several hundreds, to explore the relationship between the Big Five personality factors: E (Extraversion), A (Agreeableness), C (Conscientiousness), N (Neuroticism), and O (Openness) [17-18, 23-25]. These studies have indicated minimal or inconsistent differences in these personality factors. Conversely, when examining individual items considered as “blood type traits,” studies have consistently shown statistically significant differences [16-17, 26-28]. This can be attributed to the limited impact of blood type on a few “micro” results: several single-question items comprising the personality factors. As each personality factor represents a “macro” result, a statistical integration of numerous these micro results, many differences related to blood type are expected to diminish to insignificance. This parallels the mechanisms observed in genes influencing traits like height, weight or education [9-11], as well as hair color [29].

Since 2000, several studies have been published to explore the relationship between biological factors [30-31], and Japanese psychiatric researchers reported that the TCI personality test [32-33], based on a 7-dimensional model of temperament and personality consisting of 240 questions, found a relationship as predicted by blood type [34]. In 2022, several papers on ABO blood type and gut microbiota were published [35-37]. In particular, differences in the abundance of bacteria have been shown to have a causal relationship with depression and mental health. In 2022, another study has also been published on ABO blood typing of more than 1 million pregnant Chinese females, using a multidisciplinary approach called assortative mating, in which male-female pairs are more likely to have the same genotype or the same blood type [38].

### 1.4 Direction of This Study

As aforementioned, a multitude of genes are implicated in macro-level phenomena such as height, weight or education [9-11]. If we assume that the properties of the 5 factors of the Big Five personality tests are similar to those of height, it becomes possible, at least theoretically, to consider these 2 mechanisms to be similar. It should be noted that the Big Five personality test ultimately condenses the vast pool of potential question items to less than a few hundred, effectively truncating items exhibiting relatively minute differences. Should genes act independently on individual micro-level items with weak or modest effects, it follows that a significant portion of question items varying by blood type would be largely eliminated during the final construction of the personality test, resulting in substantially diminished differences in macro-level personality factors. This aligns with prior studies that have failed to yield consistence and mirrors the mechanism by which multiple genes affect over attributes such as height, weight or education.

While rigorous testing of this hypothesis requires an extensive personality and genomic data, “blood type” offers a partial remedy to the situation. Notably, the blood type factor demonstrates a more favorable landscape due to the fact that over 90% of the Japanese population knows their blood types, and if absolute precision is not demanded, an Internet survey would suffice for accomplishing the task. This blood type scenario in Japan extends to other East Asian countries, such as Korea and Taiwan, owing to the presence of conscription systems so that a substantial portion of the population know their blood types. Hence, this study aims to conduct an international comparison between Japan and Korea, while employing standardized versions of the Big Five personality tests to facilitate more precise and reliable outcomes through survey analysis.

### 1.5 Analysis Using AI

It has been observed that personality demonstrates intricate interactions with genetic factors, such as gender or age, and these interactions are not necessarily linear [39-41]. However, previous personality studies have often presumed that the influence of blood type remains constant regardless of gender or age. Statistical methods employed in the design of personality tests (e.g., principal components analysis and correlation analysis) inherently assume linear associations among variables, lacking a theoretical assurance of accurately analyzing real-world, nonlinear data.

Artificial intelligence (AI) or machine learning can be employed to address these complexities. Recent AI-driven pilot research might have corroborated at least some of the aforementioned mechanisms [16]. AI possesses the capacity to theoretically handle nonlinear models. Hence, in this study, we selected to employ AI to predict blood type using the items in the questionnaire. This predictive task exemplifies a classic illustration of “supervised learning,” which does not necessitate a specialized mathematical model and can be effectively managed by contemporary AI cloud systems in common use. If the aforementioned supposition—that genes exert independent effects on each individual question item, with small impact, and that these effects may appear less in personality tests with a limited number of items, we expect that using the blood type traits to predict blood type will provide better accuracy than the 5 factor prediction of the Big Five personality tests.

## 2. Methods

### 2.1 Participants and Question Items

Due to the extensive sample size required for AI, we chose to employ crowdsourcing as a means of data collection. For the purpose of conducting the Big Five personality tests, we utilized the TIPI (10 items) [42-44] and the BFI (44 items in version 1, and 60 items in version 2) [45-48], both of which have undergone standardization in many countries and languages. In the case of the BFI, we utilized the 60-item version 2 of the BFI-2J [47] for the Japanese participants. However, since there is currently no official version 2 available for the Korean language, we employed the version 1, BFI-K [48], comprising of 44 items.

The sample size for the Japanese edition amounted to 2,000 individuals, where both Japanese males and females aged 20 to 59 years were administered 2 Big Five tests: the TIPI-J (Japanese edition of the TIPI) and the BFI-2J (Japanese edition of the BFI version 2). The sample allocation was conducted in equal proportions across gender, and age groups in 10-year increments (Table 1). Each question item was evaluated on a scale ranging from 1 to 5, with higher values indicating greater applicability. It should be noted that while the original TIPI employs a 7-point scale, the Korean edition employed a different crowdsourcing system that only supported a 5-point scale, we used a 5-point scale for consistency. For the assessment of blood type, a total of 8 items were utilized, with 2 items assigned to each of the 4 blood types (A, B, O, and AB). Participants were asked to rate the degree to which they perceived their personality to match their blood type trait items on a scale ranging from 1 to 5, with higher values indicating greater applicability. Furthermore, participants were requested to indicate their level of belief and knowledge of the relationship between blood type and personality on a scale ranging from 1 to 4, with higher values indicating greater belief or knowledge.

**Table 1.** Datasets Used.

| Dataset | Sample Size | Male | Female | Mean Age (SD) |
| --- | --- | --- | --- | --- |
| Japanese | 1,827 | 892 | 935 | 40.3 (11.2) |
| Korean | 482 | 247 | 235 | 39.9 (10.6) |

In the Korean edition, which employed a different crowdsourcing service, the sample size was set at 500 respondents, while employing the same methodology as the Japanese edition (Table 1). As aforementioned, BFI version 2 has not yet been standardized, leading to the utilization of BFI-K, a Korean edition of version 1. Similarly, for the TIPI, a 5-point scale was used for convenience rather than the original 7-point scale. In the case of blood type assessment, a total of 9 items were employed, including 3 items for type A, 2 items for type B, 3 items for type O, and 1 item for type AB, as described later. Due to constraints with equal assignment of gender and age within the Korean service, these distributions were not perfectly balanced. To facilitate international comparisons between Japan and Korea, 19 common items present in versions 1 and 2 of the BFI were utilized. The data collection for both Japanese and Koreans was conducted in 2023. Blood type information of participants was acquired through self-reporting. This approach was adopted due to the widespread familiarity of blood types among individuals of Japanese and Korean populations. In the Japanese survey, a total of 97 respondents out of 2,000 (4.9% of the total) were unaware of their blood type, while 1,903 participants knew their blood type. In the Korean survey, the number of participants who lacked knowledge of their blood type was 3 out of 500 (0.6% of the total), therefore 497 respondents knew their blood type. Among these individuals, the final analysis included 1,827 Japanese participants and 482 Koreans, excluding samples that were clearly inappropriate, such as those who provided identical numerical responses for all question items in the BFI-2J or BFI-K, as well as individuals whose age fell outside the range of 20 to 59 years. To assess the sensitivity of personality, a single item was incorporated into the questionnaire to gauge participants’ interest in both their own and others’ personalities. Participants were asked to rate their level of interest on a scale ranging from 1 to 5, with higher values denoting a greater interest.

The blood type traits assessed in this study were shown in Table 2. To ensure ethical compliance, the Japanese items were derived from published academic papers [21-22]. As for the Korean items, we incorporated 9 specific items, comprising 3 items for blood type A, 2 items for blood type B, 3 items for blood type O, and 1 item for blood type AB. These items were identified based on a study by Cho et al. in 2005 [17], which revealed variations in traits across different blood types. Each question item underwent scrutiny by 2 independent companies (according to the languages) that obtained the Japanese Privacy Mark (JIS Q15001), both of which specialize in crowdsourcing. The results confirmed that there was no issue. These companies provide anonymized data to clients and obtain informed consent from respondents beforehand. Furthermore, the study obtained prior approval from the Ethics Review Committee to ensure compliance with ethical standards.

**Table 2.** Blood Types and Its Major Personality Traits.

| Blood Type | Personality Traits |  |
| --- | --- | --- |
|  | Japanese | Korean |
| A | 1. Meticulous, 2. Nervous | 1. Timid, 2. Meticulous, 3. Introvert |
| B | 1. Self-paced, 2. Self-centered | 1. Free-spirited, 2. Hot-tempered |
| O | 1. Big-hearted, 2. Laid back | 1. Active, 2. Smooth, 3. Generous (Big-hearted) |
| AB | 1. Hard to be understood, 2. Dual personality | 1. Unique |

Data analysis was conducted using jamovi 2.3.18 statistical software, with a significance level set at α = 0.05. The distribution of blood types in the Japanese dataset consisted of 698 individuals with type A, 404 individuals with type B, 546 individuals with type O, and 179 individuals with type AB, closely mirroring the Japanese population average [49]. The distribution in the Korean dataset consisted of 151 individuals with type A, 147 individuals with type B, 137 individuals with type O, and 47 individuals with type AB. These figures closely align with the results obtained from the 2021 survey conducted by the Korean Military Manpower Administration [50]. Additionally, effect size calculations were performed [51]. For multiple comparisons, Holm’s method was employed.

### 2.2 Analytical Strategy

We adhere to established methods of psychological personality assessment, deliberately selecting traits (Table 2) that were most suitable for eliciting discernible differences. These traits were carefully chosen based on their consistency with previous academic studies, their demonstration of significant differences in those studies, their mean values being close to 50%, their lack of extreme values, and their consistency with previous surveys conducted in related studies. As previously mentioned, due to the limited size of the Korean participants obtained in this study (500 individuals), employing AI for blood type prediction through machine learning proved unfeasible. Consequently, AI-based blood type prediction was conducted only for the Japanese participants.

We utilized the Microsoft Azure Machine Learning platform, a cloud-based AI system. All personality factors and blood type traits and were employed as training data for predicting the blood type. The entire dataset was divided into 2 subsets, with 30% allocated for training the model and the remaining 70% for predicting the blood types.

AUC was used to evaluate the predictions; AUC is a metric used to evaluate binary classification. It has a value of 0.5 if machine learning does not work (the prediction is the same as a random result), and a value of 1 for a perfectly correct prediction. For multiple classifications with 4 values, such as blood type, the weighted AUC is used instead of the standard AUC. Similarly, the value of the weighted AUC is 0.5 if the prediction is the same as the random result and 1 if it is a perfect prediction.

## 3. Results

### 3.1 Analysis 1: Big Five Personality Tests

As for the Japanese Big Five personality tests (TIPI-J and BFI-2J), no factor was statistically significant at *p* < 0.05, or after the Holm’s correction (Table 3).

**Table 3.** Japanese ANOVA for the Big Five Personality Factors and Blood Type.

| Factors | Mean scores by blood type | | | | *F* | $\eta^2$ | *p* |
| --- | --- | --- | --- | --- | --- | --- | --- |
|  | A<br>(N=698) | B<br>(N=404) | O<br>(N=546) | AB<br>(N=179) |  |  |  |
| TIPI E | 5.050 | 5.134 | 5.179 | 5.134 | 0.562 | 0.001 | 0.640 |
| TIPI A | 6.974 | 6.874 | 6.828 | 6.810 | 1.137 | 0.002 | 0.333 |
| TIPI C | 5.742 | 5.708 | 5.802 | 5.849 | 0.395 | 0.001 | 0.756 |
| TIPI N | 6.423 | 6.364 | 6.264 | 6.391 | 0.863 | 0.001 | 0.460 |
| TIPI O | 5.371 | 5.391 | 5.432 | 5.542 | 0.559 | 0.001 | 0.642 |
| BFI E | 2.561 | 2.613 | 2.602 | 2.608 | 0.716 | 0.001 | 0.542 |
| BFI A | 3.172 | 3.175 | 3.200 | 3.152 | 0.558 | 0.001 | 0.643 |
| BFI C | 3.098 | 3.034 | 3.060 | 3.124 | 1.491 | 0.002 | 0.215 |
| BFI N | 3.154 | 3.061 | 3.094 | 3.131 | 1.855 | 0.003 | 0.135 |
| BFI O | 2.956 | 2.945 | 2.943 | 3.002 | 0.531 | 0.001 | 0.661 |

Note. $p < 0.05$ are highlighted in bold. \* $p < 0.05$ after the Holm's correction.

On the Japanese question item “Do you think blood type and personality are related?”, 7.6% of the respondents answered “related a lot,” 22.6% answered “related somewhat,” 30.3% answered “related a little,” 22.8% answered “not related at all,” and 16.7% answered “I don’t know.” On question item “Do you know the traits and compatibilities of blood types?”, 4.1% answered “I know a lot,” 20.0% answered “I know some,” 46.1% answered “I know a little,” and 29.8% answered “I don’t know at all.”

As for the Korean Big Five personality tests (TIPI-K and BFI-K), 2 factors, TIPI E and BFI E, were statistically significant at *p* < 0.05, but no factor was significant after the Holm’s correction (Table 4).

**Table 4.** Korean ANOVA for the Big Five Personality Factors and Blood Type.

| Factors | Mean scores by blood type | | | | *F* | $\eta^2$ | *p* |
| --- | --- | --- | --- | --- | --- | --- | --- |
|  | A<br>(N=151) | B<br>(N=147) | O<br>(N=137) | AB<br>(N=47) |  |  |  |
| TIPI E | 4.881 | 5.49 | 5.474 | 5.426 | 3.558 | 0.022 | **0.014** |
| TIPI A | 6.934 | 6.782 | 6.774 | 6.362 | 1.982 | 0.012 | 0.116 |
| TIPI C | 7.066 | 6.939 | 7.117 | 6.979 | 0.338 | 0.002 | 0.798 |
| TIPI N | 5.940 | 6.000 | 5.818 | 5.745 | 0.461 | 0.003 | 0.710 |
| TIPI O | 6.113 | 6.014 | 6.380 | 6.106 | 1.334 | 0.008 | 0.263 |
| BFI E | 2.849 | 3.006 | 2.982 | 2.907 | 2.818 | 0.017 | **0.039** |
| BFI A | 3.502 | 3.424 | 3.539 | 3.414 | 1.450 | 0.009 | 0.228 |
| BFI C | 3.508 | 3.457 | 3.588 | 3.463 | 1.263 | 0.008 | 0.286 |
| BFI N | 2.985 | 2.918 | 2.903 | 2.899 | 0.533 | 0.003 | 0.660 |
| BFI O | 3.141 | 3.171 | 3.280 | 3.130 | 1.629 | 0.010 | 0.182 |

Note. $p < 0.05$ are highlighted in bold. \* $p < 0.05$ after the Holm's correction.

On the Korean question item “Do you think blood type and personality are related?”, 12.2% of the respondents answered “related a lot,” 40.6% answered “related somewhat,” 23.0% answered “related a little,” 16.8% answered “not related at all,” and 7.4% answered “I don’t know.” On question item “Do you know the traits and compatibilities of blood types?”, 3.0% answered “I know a lot,” 25.2% answered “I know some,” 46.4% answered “I know a little,” and 25.4% answered “I don’t know at all.”

### 3.2 Analysis 2: Blood Type Traits

As for the blood type traits for the Japanese edition, all items were statistically significant at *p* < 0.05, and after the Holm’s correction. The magnitude of *η*^*2*^ was at most 0.033, and the effect size was small. All traits of the blood type with the highest score matched the respondents’ blood type traits (Table 5).

**Table 5.** Japanese ANOVA for the Blood Type Traits.

| Blood type traits | Mean scores by blood type | | | | *F* | $\eta^2$ | *p* |
| --- | --- | --- | --- | --- | --- | --- | --- |
|  | A<br>(N=698) | B<br>(N=404) | O<br>(N=546) | AB<br>(N=179) |  |  |  |
| A (2 traits) | 6.746 | 6.131 | 6.223 | 6.514 | 16.663 | 0.027 | < .001* |
| B (2 traits) | 6.395 | 6.745 | 6.372 | 6.469 | 5.844 | 0.010 | < .001* |
| O (2 traits) | 6.007 | 6.134 | 6.544 | 5.872 | 17.663 | 0.028 | < .001* |
| AB (2 traits) | 5.470 | 5.535 | 5.449 | 6.453 | 20.518 | 0.033 | < .001* |

Note. Scores for multiple trait items are totaled. Highest scores that match with blood type traits are shaded yellow, and $p < 0.05$ are highlighted in bold. \* $p < 0.05$ after the Holm's correction.

As for the blood type traits for the Korean edition, all items except AB were statistically significant at *p* < 0.05, and after the Holm’s correction. The magnitude of *η*^*2*^ was at most 0.030, and the effect size was small. All traits of the blood type with the highest score matched the respondents’ blood type traits (Table 6).

**Table 6.** Korean ANOVA for the Blood Type Traits.

| Blood type trait(s) | Mean scores by blood type | | | | *F* | $\eta^2$ | *p* |
| --- | --- | --- | --- | --- | --- | --- | --- |
|  | A<br>(N=151) | B<br>(N=147) | O<br>(N=137) | AB<br>(N=47) |  |  |  |
| A (3 traits) | 10.695 | 9.986 | 9.861 | 10.213 | 4.761 | 0.029 | **0.003\*** |
| B (2 traits) | 5.391 | 6.075 | 5.796 | 5.468 | 4.925 | 0.030 | **0.002\*** |
| O (3 traits) | 9.728 | 10.007 | 10.431 | 9.766 | 3.945 | 0.024 | **0.008\*** |
| AB (1 trait) | 2.940 | 2.966 | 2.949 | 3.128 | 0.430 | 0.003 | 0.732 |

Note. Scores for multiple trait items are totaled. Highest scores that match with blood type traits are shaded yellow, and $p < 0.05$ are highlighted in bold. \* $p < 0.05$ after the Holm's correction.

### 3.3 Analysis 3: Blood Type Predictions Using AI

On Japanese edition, AUCs which indicate the accuracy of predictions were, in descending order, 0.603 for blood type traits (8 items), 0.564 for the BFI (60-item Big Five personality test), and 0.515 for the TIPI (10-item Big Five personality test). In a comparison of the Big Five personality tests, the BFI-2J, which has a larger number of items, was more accurate (Table 7).

**Table 7.** Result of Blood Type Predictions Using AI.

| Data | Weighted AUC | N | Algorithm |
| --- | --- | --- | --- |
| Japanese BFI (5 factors with 60 items) | 0.564 | 1,827 | VotingEnsemble |
| Japanese TIPI (5 factors with 10 items) | 0.515 | 1,827 | StandardScalerWrapper |
| Japanese blood type traits (8 items) | 0.603 | 1,827 | VotingEnsemble |

## 4. Discussions

### 4.1 Consistency with Previous Studies

None of the differences in the scores of the 5 personality factors of the TIPI and BFI, respectively, by blood type in the Big Five personality tests was found to be statistically significant after applying Holm’s correction (Tables 3-4). Conversely, all blood type traits displayed statistically significant differences after applying Holm’s correction (Tables 5-6), with the exception of the Korean type AB, which had the fewest participants (47 individuals). All blood type scores consisting of the 8 Japanese and 9 Korean items assessing blood type traits were highest for the corresponding blood types. Thus, it can be inferred that nearly all the surveyed items related to blood type traits displayed the highest scores for the corresponding blood types, clearly indicating a relationship between blood type and personality.

When comparing the *p* values of the Big Five personality tests (TIPI with 10 items, and BFI with 60 items in the Japanese version 2 and 44 items in the Korean version 1), which differ in the number of question items, it was observed that the BFI, with its larger item pool, yielded lower *p* values. This suggests that differences by blood type are more likely to appear in a test of larger number of items (Table 8). Additionally, the Japanese AUC value of 0.564 for the BFI was higher than the AUC value of 0.515 for the TIPI, indicating a similar trend (Table 7).

**Table 8.**
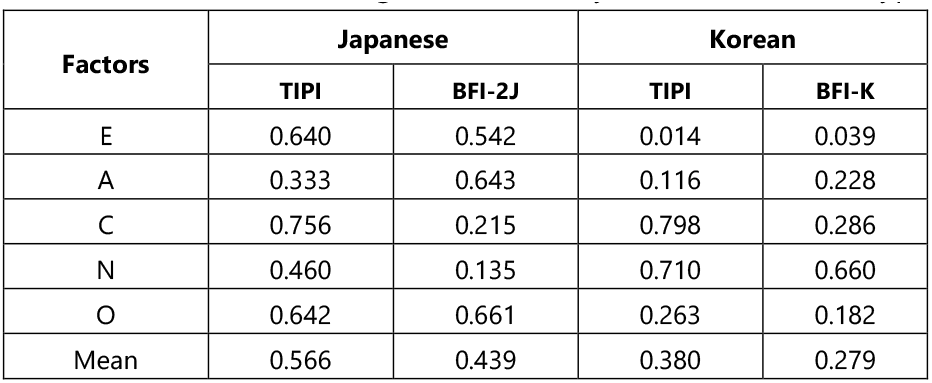
ANOVA for the Big Five Personality Factors and Blood Type.

| Factors | Japanese |  | Korean |  |
| --- | --- | --- | --- | --- |
|  | TIPI | BFI-2J | TIPI | BFI-K |
| E | 0.640 | 0.542 | 0.014 | 0.039 |
| A | 0.333 | 0.643 | 0.116 | 0.228 |
| C | 0.756 | 0.215 | 0.798 | 0.286 |
| N | 0.460 | 0.135 | 0.710 | 0.660 |
| O | 0.642 | 0.661 | 0.263 | 0.182 |
| Mean | 0.566 | 0.439 | 0.380 | 0.279 |

Items 9 (factor N) and 18 (factor C) in the Japanese BFI showed statistical significance. These items include the type A traits “optimistic (reversal item) “ and “meticulous,” respectively. Factors N and C, which include these items exhibited lower *p* values compared to the other factors (Table 3), and both factors demonstrated significant differences in individual items (Table 9). These findings support the anticipated expectation that items with minimal differences are truncated, and tests with a larger number of items are more likely to detect differences in blood type.

**Table 9.**
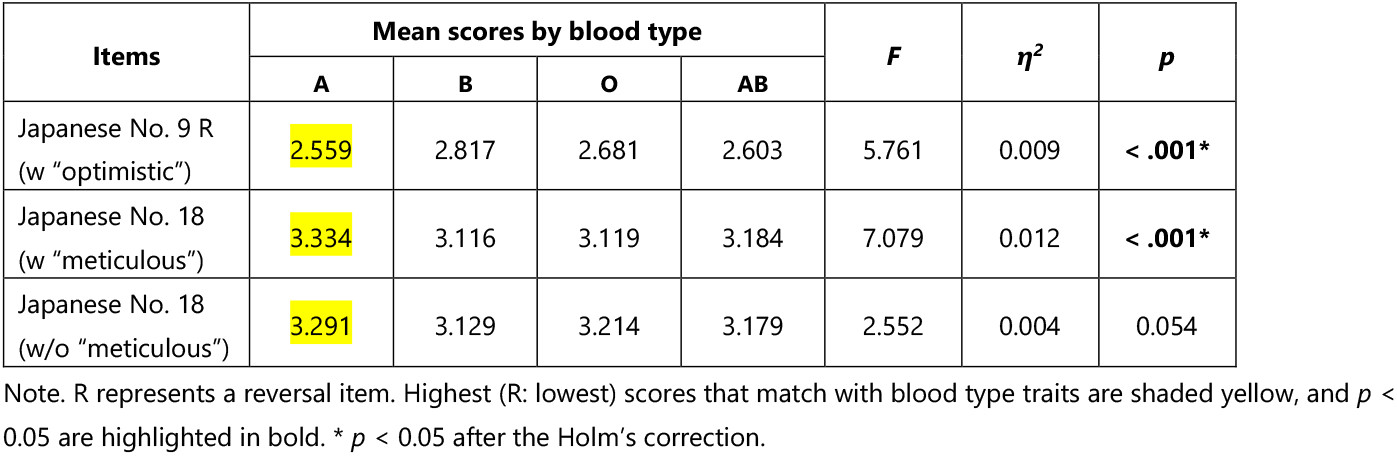
Japanese ANOVA for the BFI and Blood Type.

To further validate this phenomenon, we set the scores of items 9 and 18 in the Japanese BFI to 3 (the center value of 5-point scale), then conducted AI-based blood type prediction in a manner consistent with Analysis 3. The results revealed a decrease in the AUC value from 0.564 to 0.554. Moreover, using a provisional translation of the Japanese edition of the BFI that excluded the term “meticulous” [52], and examining the same participants as in the primary survey, the result showed no longer significant, and after applying Holm’s correction (Table 9). Hence, it becomes evident that the significance of item 18 after Holm’s correction in the Japanese edition of the BFI was primarily influenced by the inclusion of the blood type trait “meticulous,” highlighting the impact of these 2 specific items. Consequently, the influence of blood type does not display uniformly across all personality factors, but rather in a very limited number of question items. It should be noted that the Korean edition of the BFI lacks the same items as in the Japanese edition, making a direct comparison with the Japanese edition unfeasible. These results also revealed that even if the Big Five personality tests exhibit significant differences by blood type in individual items, such differences may not be reflected in the scores of the 5 personality factors.

Furthermore, when comparing the Japanese and Korean results, instances were observed where different outcomes emerged for the same items. In the Korean BFI, there is no question item that includes the Korean edition of the Type A trait “introversion,” such as item 1 of the Korean TIPI [44]. Among the items indicating extraversion in the Korean BFI edition, the results for question item 1 and question item 31 were found to be significant (Table 10). The corresponding question items in Japanese edition are No. 46 and No. 31, respectively, but neither of them displayed statistical significance. The same applied to the TIPI. Meanwhile, in the 2021 USA study, introversion was considered a type B trait, the opposite of the Korean trait [31]. These results suggest that sociocultural or linguistic structure could potentially affect the expression of personality by blood type.

**Table 10.** ANOVA for the BFI and Blood Type.

| Items | Mean scores by blood type | | | | $F$ | $\eta^2$ | $p$ |
| --- | --- | --- | --- | --- | --- | --- | --- |
|  | A | B | O | AB |  |  |  |
| Korean No. 1 | 2.662 | 2.905 | 2.555 | 2.489 | 3.643 | 0.022 | **0.013** |
| Japanese No. 46<br>(equals Korean No. 1) | 2.675 | 2.738 | 2.705 | 2.698 | 0.291 | 0.000 | 0.832 |
| Korean No. 31 R | 3.589 | 3.279 | 3.241 | 3.404 | 4.696 | 0.029 | **0.003** |
| Japanese No. 31 R<br>(equals Korean No. 31) | 3.421 | 3.307 | 3.330 | 3.380 | 1.255 | 0.002 | 0.288 |

Note. R represents a reversal item. Highest (R: lowest) scores that match with blood type traits are shaded yellow, and $p < 0.05$ are highlighted in bold. \* $p < 0.05$ after the Holm's correction.

In general, the effect of blood type on personality is smaller than that of gender or age (Figures 1-2). Note that the age of the Korean dataset is excluded from the graph because the sample size was small (500 individuals) and *p* values could not be calculated for each of the 40 groups ranging in age from 20-59 years (Figure 2).

**Figure 1.**
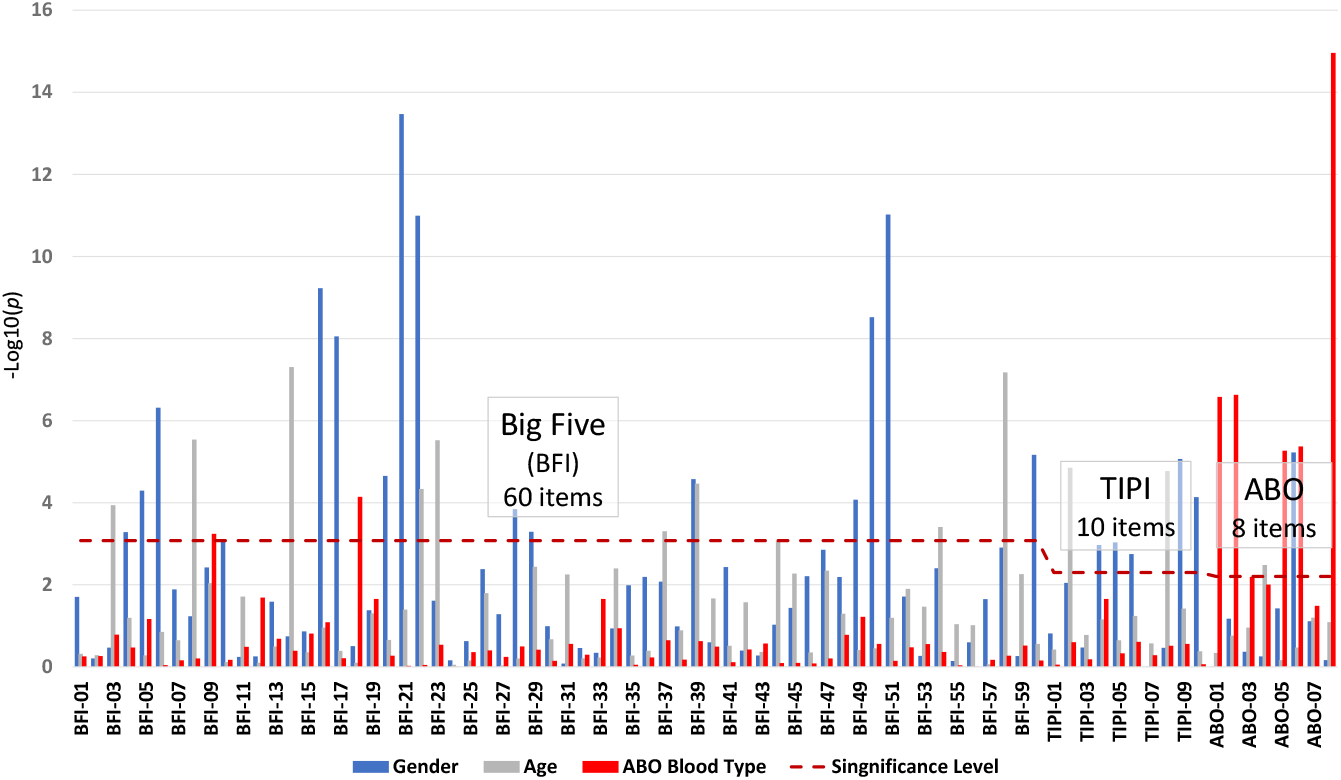
Each of Japanese Item’s *p* Value

**Figure 2.**
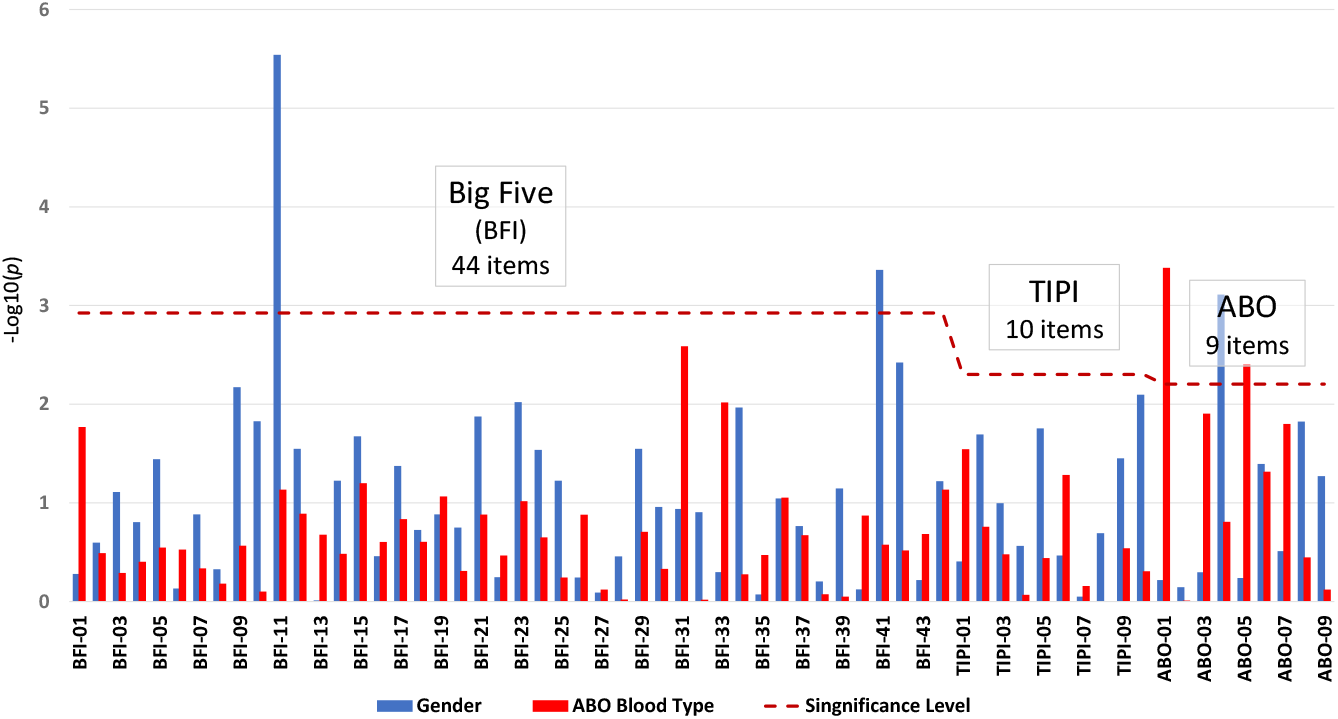
Each of Korean Item’s *p* Value

It has been reported that the Big Five personality tests also show differences in results depending on gender and/or age [39-41]. In many conventional studies, conditions other than blood type were not controlled for in many cases, because it is implicitly assumed that the effect of blood type is greater than gender, age, or other factors. Therefore, when comparing results from conventional studies, it may not be unnatural to find inconsistent results from the Big Five personality tests among them, as the effects of gender or age are likely to outweigh the effects of blood type.

Both *η*^*2*^ and *R*^*2*^ are values that indicate the magnitude of the effect on the overall variance. The *η*^*2*^ values for several traits of ABO blood type in this study were in the upper 2% range or greater (Tables 5-6), which exceed the *R*^*2*^ value in the lower 2% range or smaller for personality factors with a total gene count exceeding 400 [13]. This suggests a challenge for the Big Five personality test. Therefore, it can be deduced that differences in the 5 personality factors of this test related to blood type are relatively limited. Conversely, the differences in blood type that might have surfaced in certain items of this personality test are likely to disappear when incorporated into the 5 personality factors, owing to differences in gender, age, personality sensitivity, social environment, or other influencing factors. The fact that the same traits show differences depending on the gender (Table 11) and the age of the respondent is also consistent with the general idea that there are individual differences in gene expression.

**Table 11.**
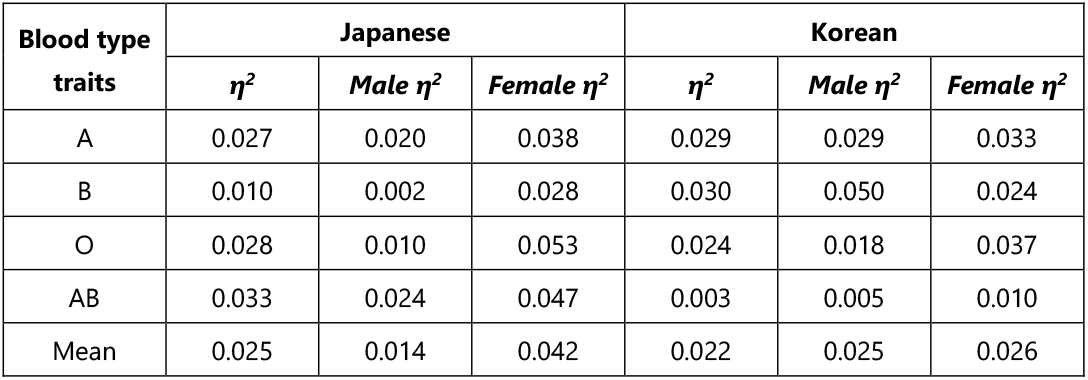
ANOVA for the Blood Type Traits by Gender in This Study.

It is noteworthy that some researchers have contended that personality differences by blood type are not real, but rather pseudo-correlations resulting from the “self-fulfillment phenomenon.” According to the findings of the 2 personality tests in this study among Korean participants, the trait of Extroversion demonstrated statistical significance at a level of *p* < 0.05 for each test. Notably, in Table 4, it can be observed that type B individuals exhibited the highest scores. Although the significance was not observed after the Holm’s correction, the agreement between the two sets of results suggests that Korean individuals with type B blood were extrovert, as anticipated. Furthermore, an earlier study conducted by Watanabe in 1994 revealed that the choice of responses, such as those provided in this study, was primarily influenced by direct association with the individual’s own personality, as opposed to similarity with impressions or archetypal traits [22].

### 4.2 Personality Sensitivity

The impact of gender and age on personality test scores is generally greater than that of blood type, as illustrated in Figures 1-2. However, based on the empirical data, it becomes apparent that “personality sensitivity,” or the responsiveness to an individual’s personality, has a more pronounced effect on personality test scores than gender or age. For instance, in the Japanese edition of the BFI’s N scale, the statement “I am interested in personality” displayed a positive correlation with all 12 question items (including 6 reversal items). Significant differences were observed in 11 out of the 12 items, and after Holm’s correction, differences remained significant in 10 items (Table 12). This yields a rather perplexing phenomenon: individuals who exhibit a greater interest in personality tend to score higher not only on items related to nervousness but also on items associated with its opposite. The most plausible interpretation of this finding is that the more interested individuals are in personality, the more inclined they are to perceive a personal relevance to any of the question items, regardless of their specific content. Consequently, personality sensitivity substantial influences on scores.

**Table 12.** Correlation Coefficients between BFI’s N items and Personality Sensitivity.

| Item No. | 4 | 9 | 24 | 29 | 44 | 49 |
| --- | --- | --- | --- | --- | --- | --- |
| Correlation Coefficients | 0.148 | 0.034 | 0.210 | 0.092 | 0.188 | 0.071 |
| *p* | **<.001\*** | 0.147 | **<.001\*** | **<.001\*** | **<.001\*** | **0.002\*** |
| Item No. (reversal items) | 14 | 19 | 34 | 39 | 54 | 59 |
| Correlation Coefficients | 0.076 | 0.207 | 0.106 | 0.112 | 0.064 | 0.179 |
| *p* | **0.001\*** | **<.001\*** | **<.001\*** | **<.001\*** | **0.006** | **<.001\*** |

Note. *p* < 0.05 are highlighted in bold. \* *p* < 0.05 after the Holm's correction.

Another perplexing observation emerged: an ANOVA results indicated that the value of *η*^*2*^, which signifies the impact of “interest in personality” on the openness factor in the Japanese version of the BFI, amounted to 0.181. This value implies that personality sensitivity accounts for 18.1% of the overall O scale exhibited in this personality test. Given that the total influence of genetics is estimated to approximately 50%, the 18.1% influence attributed to a single-question item suggests that the impact of this item alone includes roughly over one-third the influence exerted by genetic factors.

This also implies that the suitability of the Big Five personality test for assessing genetic influence may be limited. One contributing factor is that the Big Five personality test lacks a solid theoretical foundation and instead aims to maximize the variances of question item scores through principal component analysis [53-54]. Consequently, if personality sensitivity were to significantly impact scores on this test, its effect would be magnified. Moreover, based on the aforementioned findings, it is highly probable that the Big Five personality tests also exhibit substantial disparities in personality sensitivity. The influence of a single gene, as appeared in the scores, tends to be relatively minor, overshadowed by the impact of personality sensitivity. Thus, in order to accurately measure personality, it might be essential to consider personality sensitivity and employ robust indices and items that are less susceptible to its influence. Simultaneously, this implies that even when Big Five personality test scores are identical, the underlying personalities may not necessarily be the same. Drawing a parallel with sensory testing methods used to evaluate human taste, standardized by ISO 3972 and other standards, human senses display individual differences and subjectivity. The same principle can be applied to personality test scores, which may exhibit a similar situation. In fact, analysis of the Japanese and Korean datasets demonstrated that the influence of personality sensitivity on item scores outweighed that of gender or age (Figures 3-4). As with Figure 2, age was omitted from Figure 4 because the sample size for the Korean data set was small (500 respondents) and *p* values could not be calculated for each of the 40 age groups from 20 to 59.

**Figure 3.**
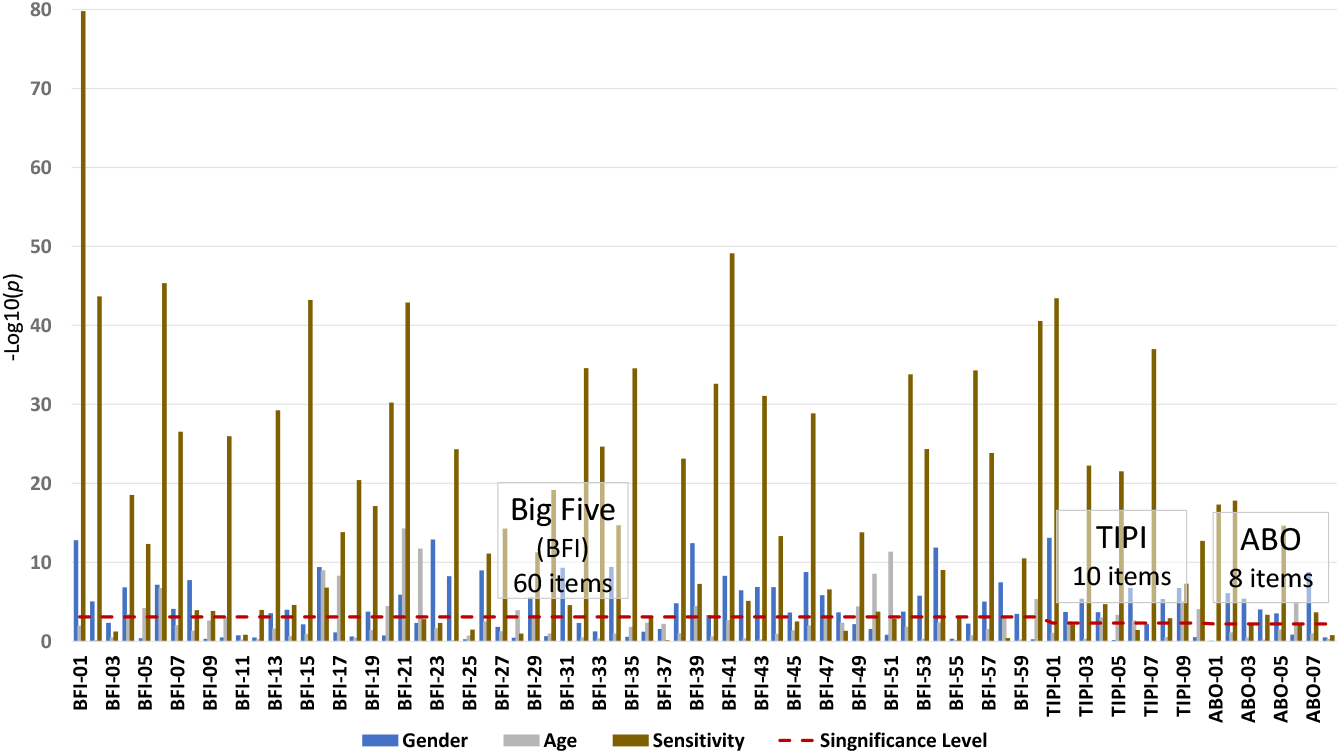
Each of Japanese Item’s *p* Value

**Figure 4.**
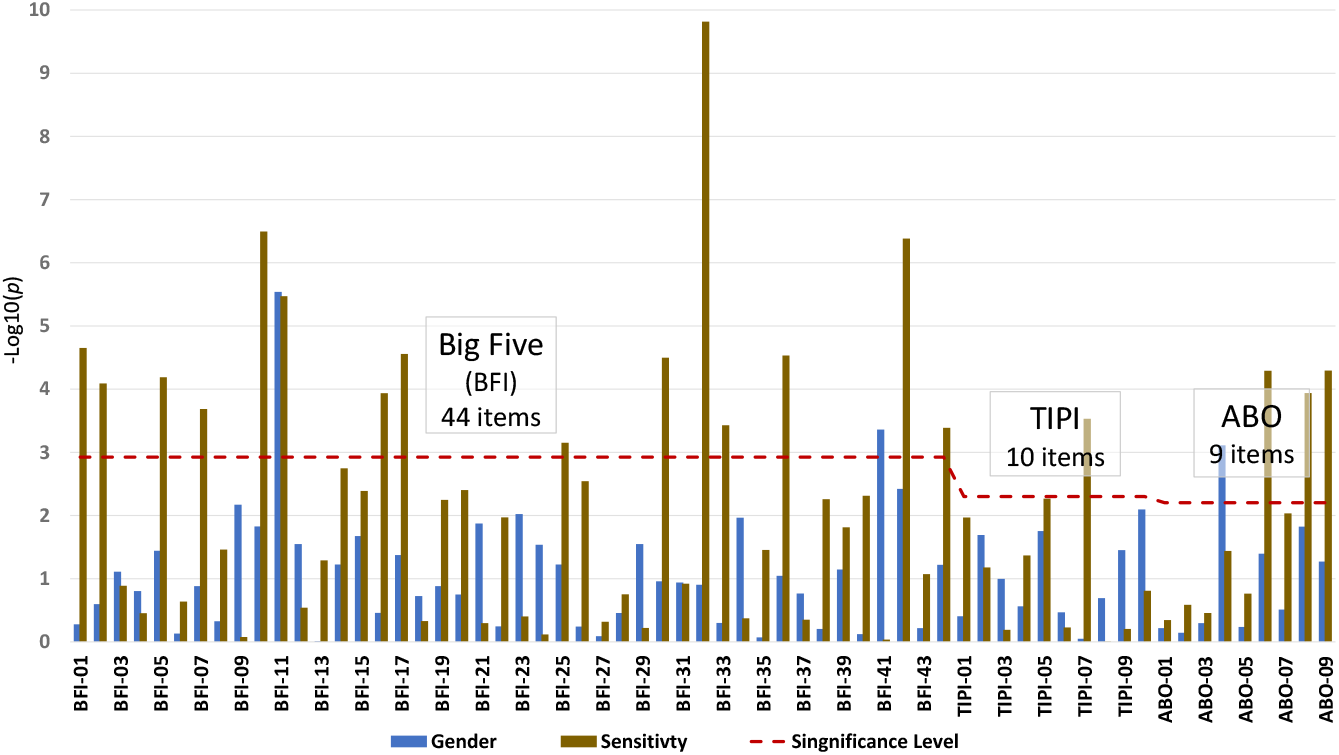
Each of Korean Item’s *p* Value

In comparison to the Big Five personality test items, the blood type traits were found to be relatively robust to personality sensitivity (Figures 1-4). This suggests that when examining the relationship between blood type and personality, blood type traits might be more suitable than the currently prevalent Big Five personality tests. These results offer valuable insights into the interpretation of personality test scores, including the possibility of Simpson’s paradox. However, given the limited number of studies on personality sensitivity [55-56], further research in this field is eagerly awaited.

### 4.3 Perspective of Evolutionary Psychology

Personality can be influenced by cultural, social, as well as genetic factors. In this study, findings from evolutionary psychology have revealed complexities that can lead to conflicting interpretations. A notable example is a series of investigations conducted by a Japanese group of Yamagishi et al [57-58].

Generally, East Asian populations, particularly the Japanese, are often perceived as collectivist, and they do exhibit collectivist behaviors in many ways. This phenomenon is commonly attributed to the longstanding practice of paddy rice cultivation in Japan, which necessitates close cooperation [59].

Due to its island geography and mountainous terrain, Japan possesses numerous small rivers compared to continental countries. Over thousands of years, paddy rice cultivation has been practiced in Japan within each narrow plain, organized according to its water system. Consequently, it is widely believed that the collectivist nature of the Japanese people is strongly influenced by the demands of paddy rice cultivation, which inherently requires close collaboration. Thus, from an evolutionary psychological standpoint, it can be argued that the Japanese population has a genetic predisposition towards collectivism.

However, Yamagishi’s group discovered through empirical data obtained from multiple experiments, that the Japanese display lower levels of trust and greater individualistic tendencies compared to individuals from the United States, China, or Taiwan. This finding is further supported by a recent comparative study on youth conducted between Japan and the United States [60-61].

The group reported that, in contrast to individuals from other countries, Japanese individuals are 1) more inclined to avoid risk and 2) significantly less likely to place trust in others [62]. Of these findings, the second observation directly contradicts the worldwide recognition of Japan as a “trusting society” with a low crime rate. They provided the following explanation for this discrepancy.

In essence, Japanese society can be described as a society where individuals establish a secure environment within a community engaged in paddy rice cultivation, characterized by strong bonds and stable relationships. To uphold this structure, external individuals are excluded, and people maintain relationships primarily with those they have known for long period. This system is upheld by the exchange of “negative” reputations within fixed peer groups. Conversely, in many developed countries today, an open and re-entrant system prevails, transcending secure and fixed relationships. The modern society is built on a general trust in others and allows for the pursuit of diverse opportunities. The modern system relies on the sharing of an individual’s “positive” reputation and necessitates the presence of a publicly upheld judicial system as a safety net to preserve this trust. Analogically, from an evolutionary psychology perspective, Choi and Bowles referred these 2 typical groups as parochial altruists (e.g. Japanese style) and tolerant nonaltruists (e.g. non-Japanese style), and showed that they are both evolutionary stable strategies (ESS) through game-theoretic analysis and agent-based simulations [63].

Later, Yamagishi et al. actually conducted an Internet auction experiment with 791 participants to examine whether this closed or open system was more economically efficient. The participants were divided into 29 groups and actually bought and sold goods with other participants in an Internet auction set up on a computer in a laboratory. The results showed that in the case of the now-common open market system, which allows for re-entry, the sales of the group that emphasized positive reputation exceeded those of the group that emphasized negative reputation. Even in real history, Maghribi merchants who had adopted a negative reputation system lost their supremacy in Mediterranean trade to Genoese merchants who had adopted a positive reputation system. This could also be one of the reasons for the current low growth of the Japanese economy. Thus, when considering from an evolutionary psychological perspective, the interpretation of the results based on conventional findings, such as the results of the Big Five personality tests, may appear contradictory at first glance, and will require careful and multifaceted analysis.

## 5. Conclusion

The expression of traits governed by a single gene is limited. Nevertheless, we observed a clear relationship between blood type and self-reported personality traits in several single-question items, aligning with previously discussed traits. Conversely, traditional personality assessments, such as the Big Five personality test, typically include several personality factors composed of multiple items. Blood type alone does not affect the personality of the individual. Rather, a multitude of factors, including gender, age or sociocultural factors, intricately shape and significantly impact personality outcomes. Without accounting for these nonlinear interactions in individual items, achieving consistent results becomes unattainable. In the context of traditional personality tests like the Big Five, the influence of blood type fluctuates, resulting in inconsistent findings unless the sample is appropriately controlled for variables such as gender, age, or other relevant conditions. The aforementioned explanation matches the theories within the field of personality psychology, as well as with other instances of genetic effects on human traits. Our discoveries propose a novel, albeit speculative, framework for understanding the influence of human genes. It should be noted that this study’s sample was confined to Japanese and Korean populations, the AI training data was limited in size and experimental, and the interactions between gender and age remains unknown. Further investigations employing more extensive and globally diverse datasets are necessary to address the true implications and refine methodologies.

## Supporting information

S1 Dataset. Raw data.

## Acknowledgements

The author expresses appreciation to Chieko Ichikawa, Director of the Human Sciences ABO Center, for her assistance, as well as to Fred Wong, co-founder of AI Hong Kong Limited, for his guidance in utilizing AI. The author also extends gratitude to Professor Qinglai Meng of Oregon State University for his feedback and recommendations.

## Supplemental File

S1 Dataset. Raw data. (XLSX)

## References

1. Zwir I, Arnedo J, Del-Val C et al. Uncovering the complex genetics of human temperament. Mol Psychiatry. 2020;25(10):2275–2294. doi:10.1038/s41380-018-0264-5

2. Ooki S. Twin Study in Physiological Anthropology. Japanese Journal of Physiological Anthropology. 2017;22(2):97–105. doi:10.20718/jjpa.22.2_97

3. Plomin R, DeFries JC, Knopik VS, Neiderhiser JM. Top 10 Replicated Findings From Behavioral Genetics. Perspect Psychol Sci. 2016;11(1):3–23. doi:10.1177/1745691615617439

4. Polderman TJ, Benyamin B, de Leeuw CA et al. Meta-analysis of the heritability of human traits based on fifty years of twin studies. Nat Genet. 2015;47(7):702–709. doi:10.1038/ng.3285

5. Bruder CE, Piotrowski A, Gijsbers AA et al. Phenotypically concordant and discordant monozygotic twins display different DNA copy-number-variation profiles. Am J Hum Genet. 2008 Mar;82(3):763–71. doi: 10.1016/j.ajhg.2007.12.011

6. Jonsson H, Magnusdottir E, Eggertsson HP et al. Differences between germline genomes of monozygotic twins. Nat Genet. 2021;53,27–34. doi:10.1038/s41588-020-00755-1

7. Dubois L, Ohm Kyvik K, Girard M et al. Genetic and Environmental Contributions to Weight, Height, and BMI from Birth to 19 Years of Age: An International Study of Over 12,000 Twin Pairs. PLOS ONE. 2012;7(2):e30153. doi:10.1371/journal.pone.0030153.

8. Shikishima C, Ando J, Ono Y et al. Registry of adolescent and young adult twins in the Tokyo area. Twin Res Hum Genet. 2006 Dec;9(6):811–6. doi: 10.1375/183242706779462769

9. Plomin R, von Stumm S. The new genetics of intelligence. Nat Rev Genet. 19;2018,148–159. doi:10.1038/nrg.2017.104

10. Okbay A, Wu Y, Wang N et al. Polygenic prediction of educational attainment within and between families from genome-wide association analyses in 3 million individuals. Nat Genet. 2022;54m437–449. doi:10.1038/s41588-022-01016-z

11. Yengo L, Vedantam S, Marouli E et al. A saturated map of common genetic variants associated with human height. Nature. 2022;610(7933):704–712. doi:10.1038/s41586-022-05275-y

12. Lo MT, Hinds DA, Tung JY et al. Genome-wide analyses for personality traits identify six genomic loci and show correlations with psychiatric disorders. Nat Genet. 2017;49(1):152–156. doi:10.1038/ng.3736

13. Hindley G, Shadrin, AA, van der Meer D. et al. Multivariate genetic analysis of personality and cognitive traits reveals abundant pleiotropy. Nat Hum Behav. 2023. doi: 10.1038/s41562-023-01630-9

14. Furukawa T. A study of temperament and blood groups. Journal of Social Psychology. 1930; 1:494–509.

15. Furukawa T. A Study of Temperament by means of Human Blood Groups. Japanese Journal of Psychology. 1927; 2(4):612–634. doi:10.4992/jjpsy.2.612

16. Kanazawa M. Pilot Analysis of Genetic Effects on Personality Test Scores with AI: ABO Blood Type in Japan. Bio Med. 2023; 15:524. doi: 0.35248/0974-8369.23.15.524.

17. Cho SH, Suh EKM, Ro YJ. Beliefs about blood types and traits and their reflections in self-reported personality. Korean Journal of Social and Personality Psychology. 2005; 19(4): 33–47. Retrieved from http://kiss.kstudy.com/thesis/thesis-view.asp?key=2498184 on June 11, 2023.

18. Wu K, Lindsted KD, Lee JW. Blood type and the five factors of personality in Asia. Pers Individ Dif. 2005; 384: 797–808. doi: 10.1016/j.paid.2004.06.004

19. Nomi M. Blood Type Humanics. 1973; Tokyo: Sankei Publishing.

20. Nomi T, Besher A. You Are Your Blood Type. 1988: New York: Pocket Books.

21. Sato T, Watanabe Y. Psychological studies on blood-typing in Japan. Japanese Psychological Review. 1992;35: 234–268. doi:10.24602/sjpr.35.2_234

22. Watanabe Y. The roles of prototype and exemplar in the formation of the “blood type stereotype”. Japanese Journal of Social Psychology. 1994;10(2):77–86. doi: 10.14966/jssp.KJ00003724631

23. Cramer KM, Imaike E. Personality, blood type, and the five-factor model. Pers Individ Dif. 2002; 32: 621–626. doi:10.1016/S0191-8869(01)00064-2

24. Shimizu T, Ishikawa M. Relationships between ABO blood types and personality: Measurement by the Five Factor Model. Kozokoseishugi-kenkyu. 2011; 5:78–91.

25. Alsadi R. Personality traits and their relationship with blood groups among of Palestinian university students. International Journal of Psychology and Behavioral Sciences. 2020;10(2):34–42. doi: 10.5923/j.ijpbs.20201002.02.

26. Yamaoka S. Discrimination and delusion of blood type personality divination. 18th Annual Meeting of Japan Society of Personality Psychology. 2009;18,11. doi: 10.24534/amjspp.18.0_11

27. Sakamoto A, Yamazaki K. Blood-typical personality stereotypes and self-fulfilling prophecy: A natural experiment with time-series data of 1978-1988. Progress in Asian Social Psychology. 2004; 4, 239–262. Seoul: Kyoyook-KwahakSa.

28. Muto C, Nagashima M, Harada J et al. A demonstrative and critical study on pseudo-science for scientific literacy construction at teacher education course. Grants-in-Aid for Scientific Research FY2011 Final Research Report (Japan). 2012. Retrieved from https://kaken.nii.ac.jp/en/grant/KAKENHI-PROJECT-22650191/

29. Hysi PG, Valdes AM, Liu F,et al. Genome-wide association meta-analysis of individuals of European ancestry identifies new loci explaining a substantial fraction of hair color variation and heritability [published correction appears in Nat Genet. 2019 Jul;51(7):1190. Nat Genet. 2018;50(5):652–656. doi:10.1038/s41588-018-0100-5

30. Hobgood DK. Personality traits of aggression-submissiveness and perfectionism associate with ABO blood groups through catecholamine activities. Med Hypotheses. 2011; 77: 294–300. doi: 10.1016/j.mehy.2011.04.039

31. Hobgood DK. ABO B gene is associated with introversion personality tendencies through linkage with dopamine beta hydroxylase gene. Med Hypotheses. 2021; 148: 110513. doi: 10.1016/j.mehy.2021.110513

32. Cloninger CR, Przybeck TR, Svrakic DM, Wetzel RD. The Temperament and Character Inventory (TCI): a guide to its development and use. St.Louis, Missouri: Center for Psychobiology of Personality Washington University; 1994.

33. Kijima N, Saito R, Takeuchi M et al. Cloninger’s seven-factor model of temperament and character and Japanese version of Temperament and Character Inventory (TCI). Kikan Seishinka Shindangaku. 1996;7: 379–399.

34. Tsuchimine S, Saruwatari J, Kaneda A, Yasui-Furukori N. ABO Blood Type and Personality traits in healthy Japanese subjects. PLOS ONE. 2015;10(5):e0126983. doi: 10.1371/journal.pone.0126983

35. Yang H, Wu J, Huang X, et al. ABO genotype alters the gut microbiota by regulating GalNAc levels in pigs. Nature. 2022;606(7913):358–367. doi:10.1038/s41586-022-04769-z

36. Qin Y, Havulinna AS, Liu Y et al. Combined effects of host genetics and diet on human gut microbiota and incident disease in a single population cohort. Nat Genet. 2022;54(2):134–142. doi:10.1038/s41588-021-00991-z

37. Lopera-Maya EA, Kurilshikov A, van der Graaf A et al. Effect of host genetics on the gut microbiome in 7,738 participants of the Dutch Microbiome Project [published correction appears in Nat Genet. 2022 Sep;54(9):1448]. Nat Genet. 2022;54(2):143–151. doi:10.1038/s41588-021-00992-y

38. Hou Y, Tang K, Wang J, Zhang H. Assortative mating on blood type: Evidence from one million Chinese pregnancies. PNAS. 2022;119(51):e2209643119. doi:10.1073/pnas.2209643119

39. Kawamoto T, Oshio A, Abe S et al. Age and gender differences of Big Five personality traits in a crosssectional Japanese sample. Japanese Journal of Developmental Psychology. 2015:26(2):107–122. doi: 10.11201/jjdp.26.107

40. Lehmann R, Denissen JJ, Allemand M, Penke L. Age and gender differences in motivational manifestations of the Big Five from age 16 to 60. Dev Psychol. 2013;49(2):365–83. doi: 10.1037/a0028277

41. Weisberg YJ, Deyoung CG, Hirsh JB. Gender Differences in Personality across the Ten Aspects of the Big Five. Front Psychol. 2011; 2:178. doi: 10.3389/fpsyg.2011.00178

42. Gosling SD, Rentfrow PJ, Swann Jr WB. A very brief measure of the Big-Five personality domains. Journal of Research in Personality. 2003;37(6):504–528. doi: 10.1016/S0092-6566(03)00046-1

43. Oshio A, Abe S, Cutrone P. Development, reliability, and validity of the Japanese version of Ten Item Personality Inventory (TIPI-J). Japanese Journal of Personality. 2012;21(1):40–52. doi: 10.2132/personality.21.40

44. Retrieved from https://gosling.psy.utexas.edu/wp-content/uploads/2014/09/TIPI-Korean.pdf on June 11, 2023.

45. Soto CJ, Oliver P, John OP. Ten facet scales for the Big Five Inventory: Convergence with NEO PI-R facets, self-peer agreement, and discriminant validity. Journal of Research in Personality. 2009;43(1):84–90. doi:10.1016/j.jrp.2008.10.002

46. Soto CJ, John OP. The next big five inventory (BFI-2): developing and assessing a hierarchical model with 15 facets to enhance bandwidth, fidelity, and predictive power. J. Pers. Soc. Psychol. 2017;113, 117–143. doi: 10.1037/pspp0000096S

47. Oshino S, Shimotsukasa T, Oshio A et al. A validation of the Japanese adaptation of the Big Five Inventory-2 (BFI-2-J). Frontiers in Psychology. 2022;13: 924351. doi: 10.3389/fpsyg.2022.924351

48. Kim SY, Kim JM, Yoo JA et al. Standardization and Validation of Big Five Inventory-Korean Version (BFIK) in Elders. Korean Journal of Biological Psychiatry. 2010;17(1):15–25. Retrieved from https://www.koreamed.org/SearchBasic.php?RID=2249786 on June 11, 2023.

49. Okubo Y. Pretransfusion testing with special reference to blood groups in Japanese (2nd ed.). 1997; Tokyo: Ishiyaku Publishers.

50. Korean Military Manpower Administration (2021). Retrieved form https://kosis.kr/statHtml/statHtml.do?orgId=144&tblId=TX_14401_A043 on June 4, 2023.

51. Cohen J. Statistical power analysis for the behavioral sciences (2nd ed.). 1988; Hillsdale, NJ: Lawrence Erlbaum.

52. Retrieved form https://www.colby.edu/academics/departments-and-programs/psychology/research-opportunities/personality-lab/the-bfi-2/ on June 11, 2023.

53. Goldberg LR. An alternative “description of personality”: the big-five factor structure. J. Pers. Soc. Psychol. 1990;59(6),1216–29. doi:10.1037/0022-3514.59.6.1216

54. Goldberg LR. The development of markers for the big-five factor structure. Psychological Assessment, 1992;4,26–42. doi: 10.1037/1040-3590.4.1.26Goldberg.

55. Aron EN, Aron A. Sensory-processing sensitivity and its relation to introversion and emotionality. J. Pers. Soc. Psychol. 1997;73(2):345–368. doi: 10.1037/0022-3514.73.2.345

56. Funahashi A. Overview of studies on individual differences in sensory sensitivity. Bulletin of graduate school, Chukyo University. 2012;11(2):29–34. Retrieved from http://n8t.cn/N34GY

57. Yamagishi T, Karen SC, Watabe M. Uncertainty, trust and commitment formation in the United States and Japan. American journal of Sociology. 1998;104,165–194. doi: 10.1111/j.1467-839X.2011.01353.x

58. Liu JH, Yamagishi T, Wang F et al. Unbalanced triangle in the social dilemma of trust: Internet studies of real-time, real money social exchange between China, Japan, and Taiwan. Unbalanced triangle in the social dilemma of trust: Internet studies of real-time, real money social exchange between China, Japan, and Taiwan. Asian Journal of Social Psychology. 2011;14(4),246–257. doi:10.1111/j.1467-839X.2011.01353.x

59. Uchida Y, Takemura K, Fukushima S et al. Farming cultivates a community-level shared culture through collective activities: Examining contextual effects with multilevel analyses. J. Pers. Soc. Psychol. 2018;116(1), 1–14. doi: 10.1037/pspa0000138

60. National Institute for Youth Education of Japan. Survey of High School Students’ Views on Mental and Physical Health (2018). Retrieved from https://www.niye.go.jp/kenkyu_houkoku/contents/detail/i/126/ on June 11, 2023.

61. Arakawa K. A Big Misconception that Japanese People “Like to be with Everyone”: Is it true that Japanese people tend to be more collectivistic than the Western people. Weekly Toyokeizai. April 21, 2021. Retrieved from https://toyokeizai.net/articles/-/422892 on June 11, 2023.

62. Yamagishi T, Yoshikai N. Net reputation society. 2009; Tokyo: NTT Publishing.

63. Choi JK, Bowles S. The coevolution of parochial altruism and war. Science. 2007;318,636–640. doi:10.1126/science.1144237

